# The GET pathway safeguards against non-imported mitochondrial protein stress

**DOI:** 10.1101/2020.06.26.173831

**Authors:** Tianyao Xiao, Viplendra P.S. Shakya, Adam L. Hughes

**Affiliations:** Department of Biochemistry, University of Utah School of Medicine, Salt Lake City, UT, 84112, USA

**Keywords:** Mitochondria, ER, protein stress, protein quality control, the GET pathway, mitochondrial carrier proteins, ER-SURF

## Abstract

Deficiencies in mitochondrial import cause the toxic accumulation of non-imported mitochondrial precursor proteins. Numerous fates for non-imported mitochondrial precursors have been identified, including proteasomal destruction, deposition into protein aggregates, and mis-targeting to other organelles. Amongst organelles, the endoplasmic reticulum (ER) has emerged as a key destination for non-imported mitochondrial proteins, but how ER-targeting of these proteins is achieved remains unclear. Here, we show that the guided entry of tail-anchored proteins (GET) complex is required for ER-targeting of endogenous mitochondrial multi-transmembrane proteins. Without a functional GET pathway, non-imported mitochondrial proteins destined for the ER are alternatively sequestered into Hsp42-dependent protein foci. The ER targeting of non-imported mitochondrial proteins by the GET complex prevents cellular toxicity and facilitates re-import of mitochondrial proteins from the ER via the recently identified ER-SURF pathway. Overall, this study outlines an important and unconventional role for the GET complex in mitigating stress associated with non-imported mitochondrial proteins.

## INTRODUCTION

Mitochondria play crucial roles in energy production, metabolite synthesis, cell immunity, and apoptosis (Friedman and Nunnari, 2014). Abnormal mitochondrial function disrupts cellular homeostasis and is tightly linked to aging and many metabolic diseases (Wallace, 2005). A major consequence of mitochondrial dysfunction is the impairment of mitochondrial protein import. The vast majority of the mitochondrial proteome, which contains over 1,000 proteins, is encoded in the nucleus and translated in the cytoplasm (Pagliarini et al., 2008). Mitochondrial precursor proteins are imported into mitochondria by translocase complexes located in the outer and inner mitochondrial membranes (OMM and IMM) through a process that depends on IMM potential (Wiedemann and Pfanner, 2017). In response to mitochondrial dysfunction, mitochondrial protein import is impaired and non-imported proteins accumulate outside of mitochondria (Boos et al., 2020; Hughes and Gottschling, 2012; Wang and Chen, 2015; Wrobel et al., 2015).

Previous studies found that non-imported mitochondrial proteins trigger proteotoxicity, termed mitochondrial precursor overaccumulation stress (mPOS) (Wang and Chen, 2015; Wrobel et al., 2015). To date, a collection of studies have shown that mPOS triggers a cascade of cellular responses that help to promote cellular survival. These responses include translational suppression and proteasomal destruction in the cytoplasm, nucleus and at the mitochondrial surface (Boos et al., 2020; Hansen et al., 2018; Itakura et al., 2016; Mårtensson et al., 2019; Shakya et al., 2020; Wang and Chen, 2015; Wrobel et al., 2015). In a recent screen to elucidate fates of non-imported mitochondrial proteins, we identified the endoplasmic reticulum (ER) as an organelle to which many non-imported mitochondrial membrane proteins were targeted (Shakya et al., 2020). This observation is consistent with other studies that have also identified alternative targeting of mitochondrial proteins to the ER under a variety of conditions (Friedman et al., 2018; Hansen et al., 2018; Vitali et al., 2018). However, it remains unclear how the proteins identified in our previous screen are targeted to the ER, and what impact this has on the cell.

Here we sought to characterize the ER-targeting pathway of non-imported mitochondrial proteins and determine its role in mitigating mitochondrial protein stress during conditions of mitochondrial impairment. We found that the guided entry of tail-anchored proteins (GET) complex, a known post-translational ER-insertion pathway for C-terminal tail-anchored (TA) proteins (Schuldiner et al., 2008), is required for targeting non-imported mitochondrial carrier proteins to the ER. Specifically, we find that Get3, the cytosolic ATPase of the GET pathway (Schuldiner et al., 2008), physically interacts with polytopic non-imported mitochondrial membrane proteins. In the absence of a functional GET pathway, ER-destined non-imported mitochondrial proteins instead localize to Hsp42-dependent cytosolic foci that associate with both mitochondria and the ER. In addition, we show that by targeting non-imported mitochondrial proteins to the ER, the GET complex prevents cellular stress and provides substrates for the ER-SURF pathway (Hansen et al., 2018) to promote re-import of ER-localized mitochondrial proteins. Thus, it appears that the GET pathway plays an important role in maintaining cellular protein homeostasis in response to mitochondrial import failure.

## RESULTS

### Non-imported mitochondrial proteins are targeted to the ER

We previously conducted a microscopy-based screen using the budding yeast GFP clone collection to study the localization and abundance of over 400 mitochondrial proteins under conditions of mitochondrial membrane depolarization induced by the ionophore trifluoromethoxy carbonyl cyanide phenylhydrazone (FCCP) (Shakya et al., 2020). Through the screen, the ER was identified as a destination for approximately 3% of the non-imported mitochondrial proteome, in agreement with prior observations of mitochondrial proteins aberrantly localizing to the ER (Hansen et al., 2018; Vitali et al., 2018). We verified the localization of eight ER-localized candidates using newly generated yeast strains in which mitochondrial proteins of interest were endogenously tagged with GFP at their C-termini, and the OMM protein Tom70 was fused to mCherry to mark mitochondria (Hughes and Gottschling, 2012; Hughes et al., 2016). Using super-resolution microscopy, we found that in untreated cells, all eight proteins localized to mitochondria as expected. Upon FCCP treatment, these proteins localized to structures characteristic of yeast ER, in addition to residual mitochondrial localization (Fig. 1A, 1B, Fig. S1A-S1C). Most of these proteins were mitochondrial membrane proteins, including both OMM proteins, e.g., Alo1 (Fig. 1A), and IMM proteins, e.g., Oac1 (Fig. 1B). ER localization of these non-imported mitochondrial proteins was confirmed by their colocalization with mCherry-tagged Sec61, a component of the ER-localized translocon (Aviram and Schuldiner, 2017; Young et al., 2012) (Fig. 1C-1E, Fig. S1D-S1F). In the presence of cycloheximide, which inhibits protein synthesis, ER localization of Alo1 and Oac1 was undetectable upon FCCP treatment (Fig. S1G, S1H), indicating only newly synthesized Alo1 and Oac1 were targeted to the ER. C-terminal FLAG-tagged Alo1 and Oac1 were also targeted to the ER upon FCCP treatment as determined by indirect immunofluorescence, similar with their GFP-tagged counterparts (Fig. S2A, S2B). Thus, ER localization of these mitochondrial proteins was not due to their C-terminal GFP fusion. In addition to FCCP, we also used genetic tools to specifically block mitochondrial import via deletion of *TOM70* and *TOM71*. Tom70 and Tom71 reside on the OMM and facilitate the import of both Alo1 and Oac1 (Wiedemann and Pfanner, 2017). In *tom70/tom71*Δ mutants but not in wild type cells, Alo1-GFP or Oac1-GFP colocalized with Sec61-mCherry (Fig. 1F-1H). These results confirm that several mitochondrial proteins are alternatively targeted to the ER in response to either acute or constitutive mitochondrial import blockade.

**Figure 1.**
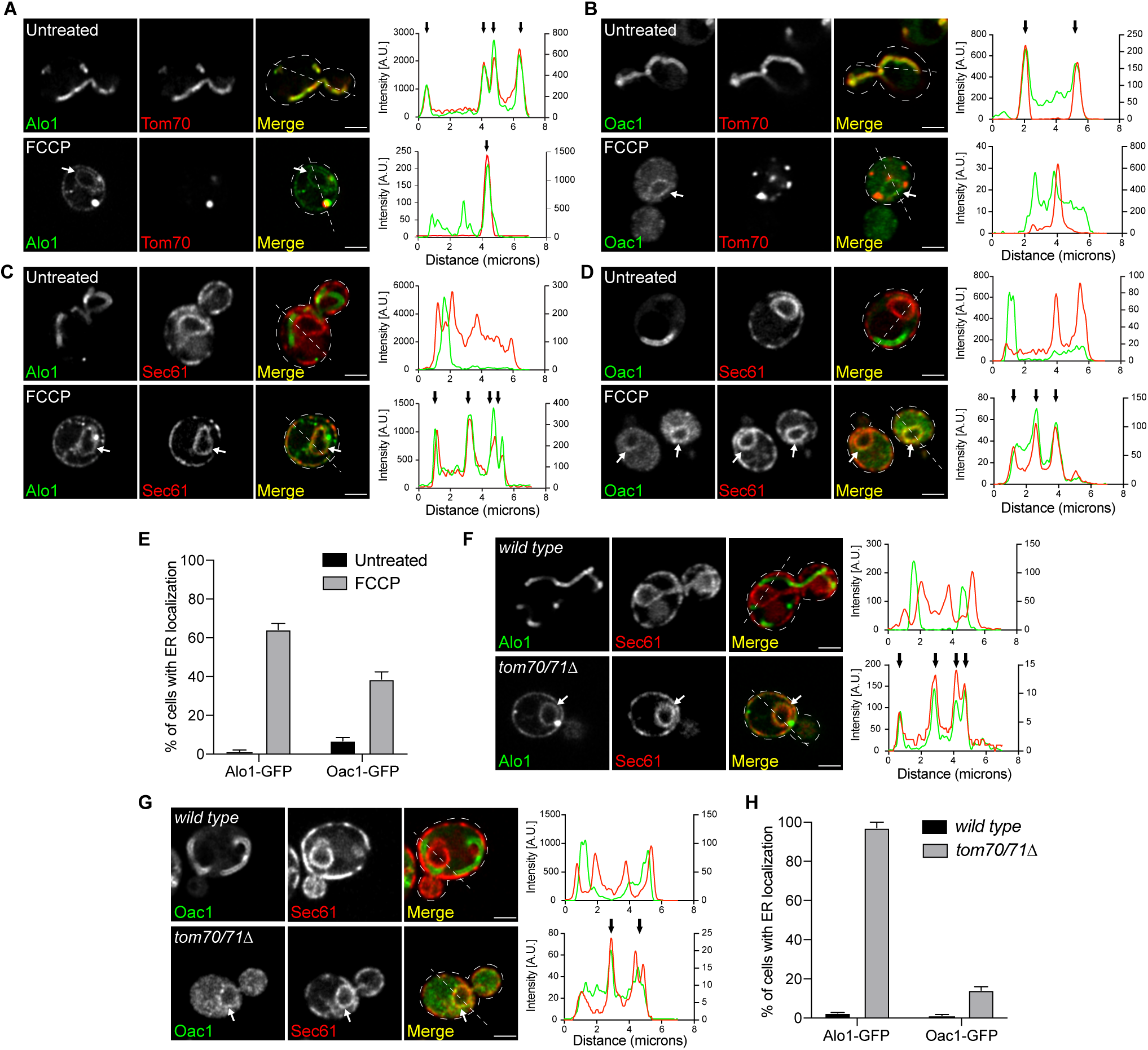
Non-imported mitochondrial proteins are targeted to the ER. (A and B) Super-resolution images and line scan analysis of yeast expressing Alo1-GFP (A) or Oac1-GFP (B) and Tom70-mCherry -/+ FCCP. (C and D) Super-resolution images and line scan analysis of yeast expressing Alo1-GFP (C) or Oac1-GFP (D) and Sec61-mCherry -/+ FCCP. (E) Quantification of cells with ER localization of Alo1- or Oac1-GFP -/+ FCCP. N > 100 cells per replicate, error bars = SEM of three replicates. (F and G) Super-resolution images and line scan analysis of *wild type* or *tom70/71*Δ expressing Alo1-GFP (F) or Oac1-GFP (G) and Sec61-mCherry. (H) Quantification of cells with ER localization of Alo1- or Oac1-GFP in wild type cells or *tom70/71*Δ mutants. N > 100 cells per replicate, error bars = SEM of three replicates. For (A-D, F and G), white arrow marks perinuclear ER. White line marks fluorescence intensity profile position. Left and right Y axis (line scan graph) correspond to GFP and mCherry fluorescence intensity respectively. Black arrow (line scan graph) marks colocalization. Images show single focal plane. Scale bar = 2 μm. See also Figure S1.

### The GET complex is required for ER-targeting of non-imported mitochondrial carrier proteins

To investigate the cellular machinery that targets non-imported mitochondrial proteins to the ER, we surveyed non-imported mitochondrial protein localization in a set of strains with deficiencies in known ER-import pathways, including the Sec61 translocon that translocates ER proteins through either the signal recognition particle (SRP)-dependent or SRP-independent pathways, the ER membrane protein complex (EMC), the SRP-independent targeting (SND) complex and the GET complex (Ast and Schuldiner, 2013; Aviram and Schuldiner, 2017; Aviram et al., 2016; Chitwood et al., 2018; Guna et al., 2018; Schuldiner et al., 2008; Shurtleff et al., 2018). Alo1 or Oac1 were endogenously tagged with GFP in mutants with deletion of either SRP-independent Sec61 translocon component *SEC72*, EMC component *EMC2*, SND complex components *SND2*, or GET pathway insertases *GET1/2* (Aviram et al., 2016; Schuldiner et al., 2008; Shurtleff et al., 2018; Wang et al., 2014a). In response to FCCP, the ER localization of Alo1 was unaffected in any of these mutants (Fig S2C, S2D). Likewise, the ER localization of Oac1 in *sec72*Δ, *emc2*Δ and *snd2*Δ upon FCCP treatment was similar to wild type (Fig S2E). In contrast, blocking the GET pathway, which normally facilitates post-translational insertion of TA proteins to the ER (Schuldiner et al., 2008), prevented FCCP-induced ER-targeting of non-imported Oac1 (Fig. 2A, 2B). Interestingly, in *get1/2*Δ mutant cells, Oac1 was sequestered in bright protein foci (Fig. 2A, 2C), consistent with previous observations that TA proteins localize to protein aggregates in GET mutants (Powis et al., 2013; Schuldiner et al., 2008). We examined additional ER-targeted non-imported mitochondrial proteins in cells lacking Get1/2 and found that the FCCP-induced ER-targeting of Mir1 and Dic1, both members of the multi-pass mitochondrial carrier protein family like Oac1 (Palmieri et al., 2006), was also dependent on the GET pathway (Fig. S3A). Similarly, Om45, an OMM protein, localized to the vacuole instead of the ER in *get1/2*Δ mutants upon FCCP treatment (Fig. S3B). In contrast, other ER-destined non-imported mitochondrial proteins still localized to the ER in GET-deficient cells when treated with FCCP (Fig. S3C, S3D), suggesting that like Alo1, their targeting is independent of the GET machinery and that multiple mechanisms exist to target non-imported proteins to the ER.

**Figure 2.**
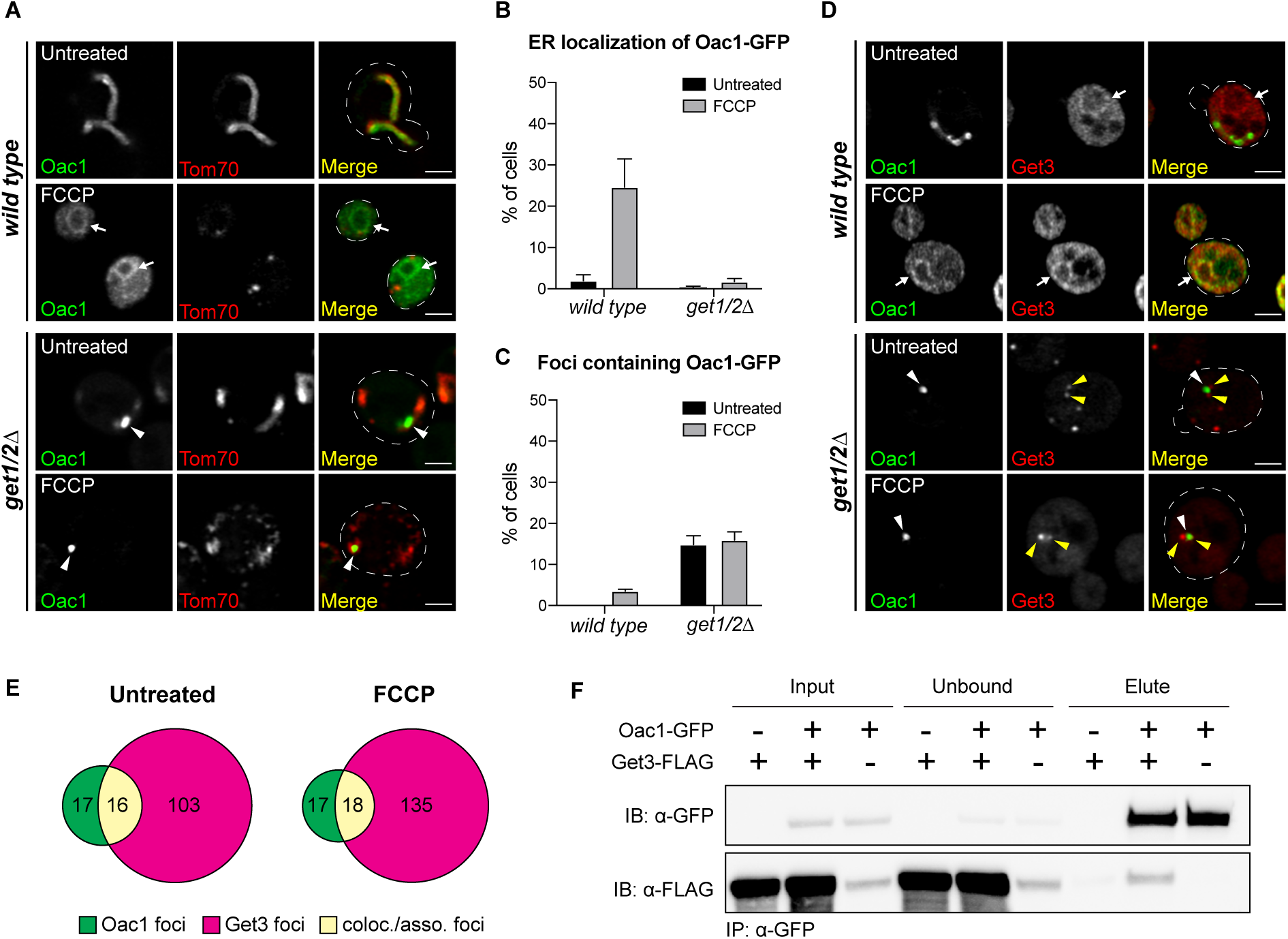
The GET complex is required for ER-targeting of non-imported mitochondrial carrier proteins. (A) Super-resolution images of wild-type or *get1/2*Δ mutant cells expressing Oac1-GFP and Tom70-mCherry -/+ FCCP. (B and C) Quantification of a showing the percentage of cells with Oac1-GFP localized to the ER (B) or protein foci (C). N > 100 cells per replicate, error bars = SEM of three replicates. (D) Super-resolution images of wild-type or *get1/2*Δ cells expressing Oac1-GFP and Get3-mCherry -/+ FCCP. (E) Quantification of (D) showing the number of foci only containing Oac1-GFP (green), Get3-mCherry (magenta) or colocalized/associated Oac1-GFP and Get3-mCherry (yellow) per 100 cells -/+ FCCP. N > 100 cells per replicate of three replicates, values are normalized to number of foci per 100 cells. (F) Western blot probing for GFP and FLAG in input, unbound or elution products of immunoprecipitated Oac1-GFP in the indicated yeast strains. White arrow marks perinuclear ER. White arrowhead marks protein foci containing Oac1-GFP. Yellow arrowheads mark protein foci containing Get3-mCherry. Images show single focal plane. Scale bar = 2 μm. See also Figure S2-S4.

To understand the involvement of other proteins of the GET complex in ER-targeting of non-imported mitochondrial proteins, we tested the requirement of upstream GET components in delivery of Oac1 to the ER, including the cytosolic ATPase Get3, which binds and recruits substrates to the ER insertases Get1/2, and the cytosolic chaperones Get4, Get5 and Sgt2, which bind and stabilize substrates to promote downstream ER-targeting by Get1/2/3 (Wang et al., 2010, 2014a). Like Get1/2, loss of Get3 also impacted targeting of Oac1 to the ER (Fig. S4A), with reduced ER-localization upon FCCP treatment (Fig. S4B) and increased number of protein foci containing Oac1 (Fig. S4C). Deletion of *GET5* also blunted ER-targeting upon FCCP treatment, but did not lead to the production of Oac1-GFP foci (Fig. S4A-S4C). Knockout of *GET4* and *SGT2*, however, had little effect (Fig. S4A-S4C). Thus, core GET components, including Get1/2/3 and partially Get5, are required for targeting mitochondrial carrier proteins to the ER, but other components of the GET pathway are dispensable.

### Cytosolic ATPase Get3 physically interacts with Oac1

To further investigate the interplay between the GET pathway components and non-imported mitochondrial carrier proteins such as Oac1, we created a strain expressing an mCherry-tagged version of Get3, the cytosolic ATPase that normally resides in the cytoplasm and recruits cytosolic GET substrates to ER-localized Get1/2 (Schuldiner et al., 2008; Wang et al., 2010). In *get1/2*Δ mutants, it has been shown that Get3 localizes to cytosolic foci containing GET substrates (Powis et al., 2013; Schuldiner et al., 2008). Like canonical TA substrates of the GET pathway (Powis et al., 2013), we found that in *get1/2*Δ mutant cells, half of the Oac1-GFP foci were colocalized or closely associated with Get3-mCherry foci, even in the absence of FCCP (Fig. 2D, 2E). Likewise, co-immunoprecipitation analysis indicated that a portion of FLAG-tagged Get3 constitutively co-purified with GFP-tagged Oac1 (Fig. 2F). This interaction persisted regardless of the nature of the epitope tags on the protein, or which of the proteins was used as the bait (Fig. S4D). In contrast, other non-ER targeted mitochondrial proteins, including Tom70, Tim50 and Por1, were not co-immunoprecipitated with Get3 (Fig. S4D). Additionally, no interaction was detected between Get3 and Alo1, a mitochondrial protein that was targeted to the ER upon impaired mitochondrial import independently of the GET complex (Fig. S4E). This indicates that Get3 selectively interacts with non-imported mitochondrial carrier proteins and promotes their ER-targeting.

### Oac1-GFP localizes to mitochondrial- and ER-associated Hsp42-dependent foci in the absence of a functional GET pathway

In cells with a non-functional GET pathway, mitochondrial carrier proteins were sequestered into protein foci (Fig 2A, 2C, Fig. S3A). To characterize these foci, we analyzed their localization in cells with fluorescently-tagged organelle markers using super-resolution microscopy. In *get1/2*Δ mutant cells, 97% of protein foci containing Oac1 were associated with mitochondria marked by Tom70 (Fig. 3A) or the ER marked by Sec61 (Fig. 3B), which is similar with previously characterized cytosolic protein aggregates (Zhou et al., 2014). To verify whether these foci corresponded to protein aggregates, we labelled Hsp42 and Hsp104, chaperones that commonly localize to cytosolic protein aggregates in yeast (Miller et al., 2015; Zhou et al., 2014), with mCherry and examined localization with Oac1-GFP foci. Interestingly, nearly all Oac1-GFP foci contained Hsp42 and Hsp104 regardless of FCCP addition (Fig. 3C-3F). Deletion of *HSP42*, but not *HSP104*, diminished the formation of Oac1-foci in *get1/2*Δ mutants (Fig. 3G-3H). Thus, without a functional GET pathway, Hsp42 mediates the sequestration of non-imported mitochondrial proteins that are normally destined to the ER.

**Figure 3.**
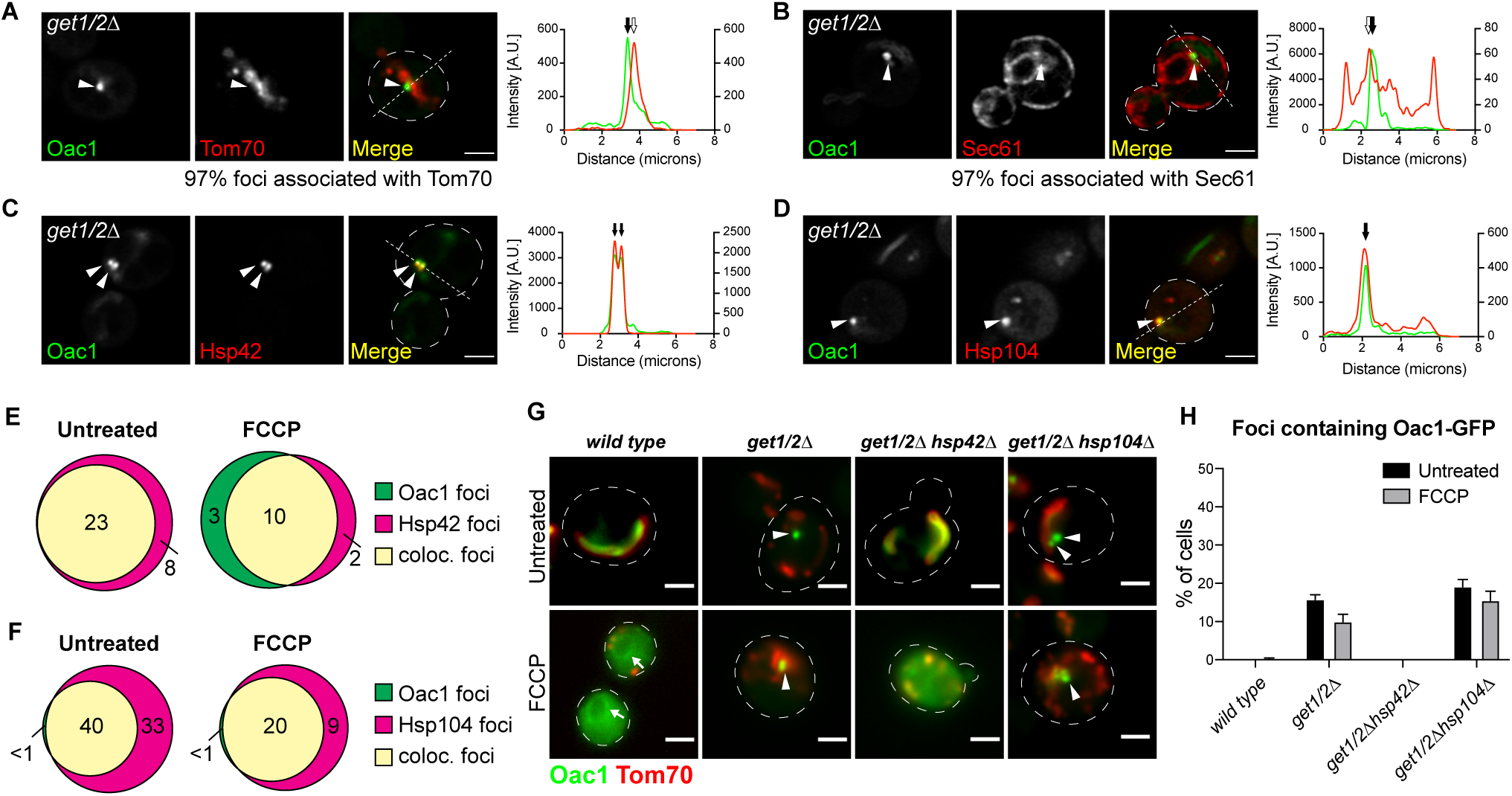
Oac1-GFP localizes to mitochondrial- and ER-associated Hsp42-dependent foci in the absence of a functional GET pathway. (A-D) Super-resolution images and line scan analysis of *get1/2*Δ mutant yeast expressing Oac1-GFP and Tom70-mCherry (A), Sec61-mCherry (B), Hsp42-mCherry (C) or Hsp104-mCherry (D). White arrowhead marks protein foci containing Oac1-GFP. White line marks fluorescence intensity profile position. Left and right Y axis (line scan graph) correspond to GFP and mCherry fluorescence intensity, respectively. Black arrow marks protein foci position and white arrow marks mitochondria (A) or ER (B) position that is associated with protein foci. For the quantification in A and B, N > 100 cells per replicate of three replicates. (E and F) Quantification of (C) and (D) respectively showing the number of foci only containing Oac1-GFP (green), Hsp42-mCherry (E) or Hsp104-mCherry (F) (magenta) or both (yellow) per 100 cells -/+ FCCP. N > 100 cells per replicate of three replicates, values are normalized to number of foci per 100 cells. (G) Widefield images of wild-type cells and the indicated mutant yeast expressing Oac1-GFP and Tom70-mCherry -/+ FCCP. White arrows mark perinuclear ER. White arrowheads mark protein foci containing Oac1-GFP. (H) Quantification of (G). N > 100 cells per replicate, error bars = SEM of three replicates. Images show single focal plane. Scale bar = 2 μm.

### GET-dependent ER targeting prevents cellular toxicity

Because non-imported mitochondrial proteins have been shown to be harmful to cells (Wang and Chen, 2015; Wrobel et al., 2015), we investigated whether the ER-targeting of non-imported mitochondrial proteins by the GET pathway prevents their toxicity. To do this, we tested the growth of GET mutants under stress of mitochondrial import failure. In comparison to wild-type cells, *get1/2*Δ cells exhibited diminished growth in the presence of FCCP (Fig. 4A), and *get3*Δ cells showed slight growth defects (Fig. 4B), in alignment with their effect on ER-targeting of mitochondrial proteins. Similarly, deletion of *GET1* or *GET2* caused fitness defects in cells lacking the mitochondrial import receptors Tom70/71 (Fig. 4C, 4D), suggesting the GET pathway mitigates cellular stress under conditions of mitochondrial import impairment.

**Figure 4.**
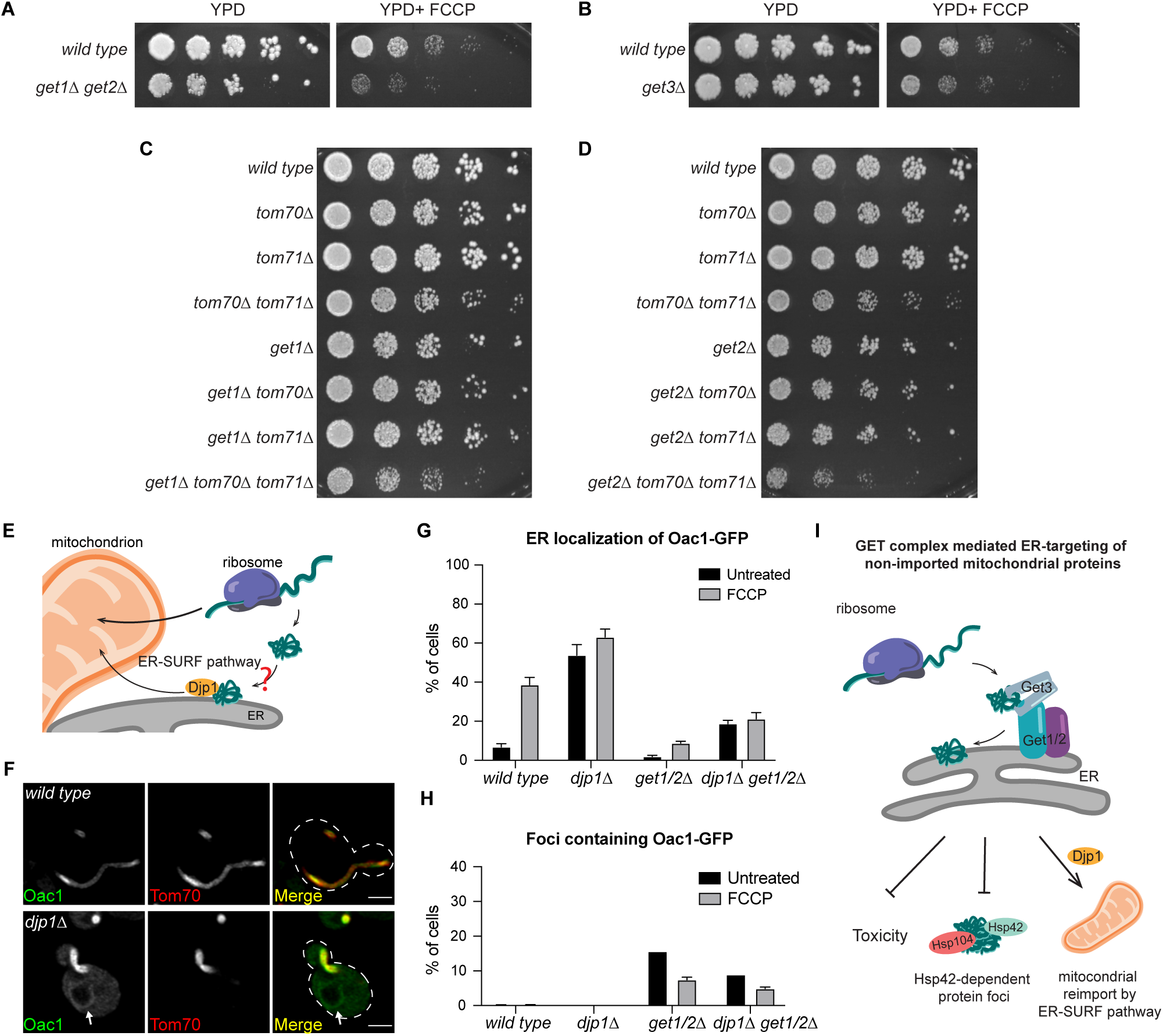
GET-dependent ER targeting prevents toxicity and provides substrates for the ER-SURF pathway. (A and B) Five-fold serial dilutions of wild-type cells and *get1/2*Δ (A) or *get3*Δ (B) mutant cells on YPD -/+ FCCP agar plates. (C and D) Five-fold serial dilutions of wild-type and the indicated mutant cells on YPD agar plates. (E) Schematic graph of the ER-SURF pathway. (F) Super-resolution images of wild-type and *djp1*Δ cells expressing Oac1-GFP and Tom70-mCherry. White arrows mark perinuclear ER. Images show single focal plane. Scale bar = 2 μm. (G and H) Quantification of the percentage of cells with Oac1-GFP localized to the ER (G) or the cytosolic foci (H) in wild-type or the indicated mutant cells. N > 100 cells per replicate, error bars = SEM of three replicates. (I) Schematic overview of the roles of GET-dependent ER targeting in preventing protein aggregation and toxicity and facilitating re-delivery of mitochondrial proteins.

### GET-dependent ER targeting provides substrates for the ER-SURF pathway

In agreement with our current findings, it was recently demonstrated that the J-protein, Djp1, shuttles ER-localized mitochondrial proteins from the ER membrane to mitochondria, promoting additional attempts of mitochondrial import (Hansen et al., 2018). A major question surrounding this pathway, termed ER-SURF, is the nature of the cellular machinery that initially targets mitochondrial proteins to the ER (Fig. 4E). With our discovery that the GET pathway facilitates ER-targeting of non-imported mitochondrial membrane proteins, we wondered whether the GET targeting system is an integral part of relaying non-imported mitochondrial precursors to the ER-SURF pathway. To test whether GET-dependent targeting of non-imported mitochondrial proteins acts upstream of Djp1, we tagged Oac1, which requires the GET complex to be localized to the ER, with GFP, in *djp1*Δ mutant cells. Interestingly, ER-localized Oac1 was observed in 55% of *djp1*Δ cells without FCCP treatment (Fig. 4F, 4G) and more than 60% with FCCP treatment (Fig. 4G). Both rates are higher than observed in wild-type cells (Fig. 4G). Importantly, the ER localization of Oac1 in *djp1*Δ cells was dramatically reduced in the absence of *GET1/2* (Fig. 4G), and protein foci containing Oac1 were present in *djp1*Δ*get1/2*Δ triple mutants (Fig. 4H). These data are consistent with a model in which the GET pathway acts upstream of the ER-SURF pathway and is required to deliver non-imported mitochondrial proteins to the ER for eventual mitochondrial re-import via Djp1.

## DISCUSSION

We previously identified the ER as a major destination of non-imported mitochondrial membrane proteins (Shakya et al., 2020). In this study, we further characterized the ER-targeting pathway of non-imported mitochondrial proteins. We show that the GET complex is required for ER-targeting of a specific group of proteins, the mitochondrial carrier proteins. With a dysfunctional GET pathway, mitochondrial membrane proteins cannot be delivered to the ER, and are instead sequestered into mitochondrion- and ER-associated cytosolic protein foci. Overall, our data support two roles for the GET pathway in quality control of ER-targeted mitochondrial proteins: mitigation of cellular stress by preventing cytosolic aggregation, and also delivery of substrates to the Djp1-dependent ER-SURF pathway for their re-import into mitochondria (Fig. 4I).

This study outlines an important and unconventional role for the GET complex in targeting multi-pass transmembrane proteins to the ER. Canonically, the GET pathway is known to insert C-terminal single-pass tail-anchored proteins to the ER (Aviram and Schuldiner, 2017; Schuldiner et al., 2008). However, our current results, along with a recent study showing that the GET complex facilitates localization of over-expressed OMM proteins to the ER (Vitali et al., 2018), suggest that the GET system is capable of handling multi-pass membrane proteins in an unknown capacity. Moving forward, it will be interesting to understand how the GET machinery structurally interacts with mitochondrial carrier proteins, including whether this interaction is direct or requires additional machinery.

Finally, our results highlight the importance of the interplay between the multitude of quality control systems that coordinately prevent toxicity induced by non-imported mitochondrial precursors. While we found that cells protect against mitochondrial protein stress through alternative ER targeting via the GET pathway, several ER-destined mitochondrial proteins analyzed here did not appear to rely on the GET pathway for their ER targeting. Identifying the systems that target these proteins will be necessary to fully understand how alternative ER delivery prevents the toxicity of non-imported mitochondrial precursors. Moreover, the degree of coordination between the many systems that mitigate the stress associated with non-imported mitochondrial precursors remains unclear. For example, while our current results suggest that GET-dependent ER targeting plays a role upstream of the ER-SURF pathway, whether the GET machinery directly communications with the ER-SURF components remains unknown. Addressing some of these questions is key to unlocking the full extent to which the non-imported mitochondrial proteome impairs cellular health during times of mitochondrial dysfunction.

## ACKNOWLEDGEMENTS

We thank members of the A.L.H. and J.M.S. laboratory for discussion and manuscript comments, and Dr. Nikolaus Pfanner for Tim50 and Tom70 antisera. We thank Janet Shaw (Utah) for contributing stipend support for T.X. Research was supported by NIH grants GM119694 (A.L.H.) and the Howard Hughes Medical Institute (J.M.S.). A.L.H. was further supported by an American Federation for Aging Research Junior Research Grant, United Mitochondrial Disease Foundation Early Career Research Grant, Searle Scholars Award, and Glenn Foundation for Medical Research Award.

## AUTHOR CONTRIBUTIONS

Conceptualization, T.X., V.P.S.S., A.L.H.; Methodology, T.X., V.P.S.S., A.L.H.; Formal Analysis, T.X.; Investigation, T.X.; Writing T.X. and A.L.H.; Visualization, T.X.; Supervision, A.L.H.; Funding Acquisition, A.L.H.

## FIGURE LEGENDS

**Figure S1.**
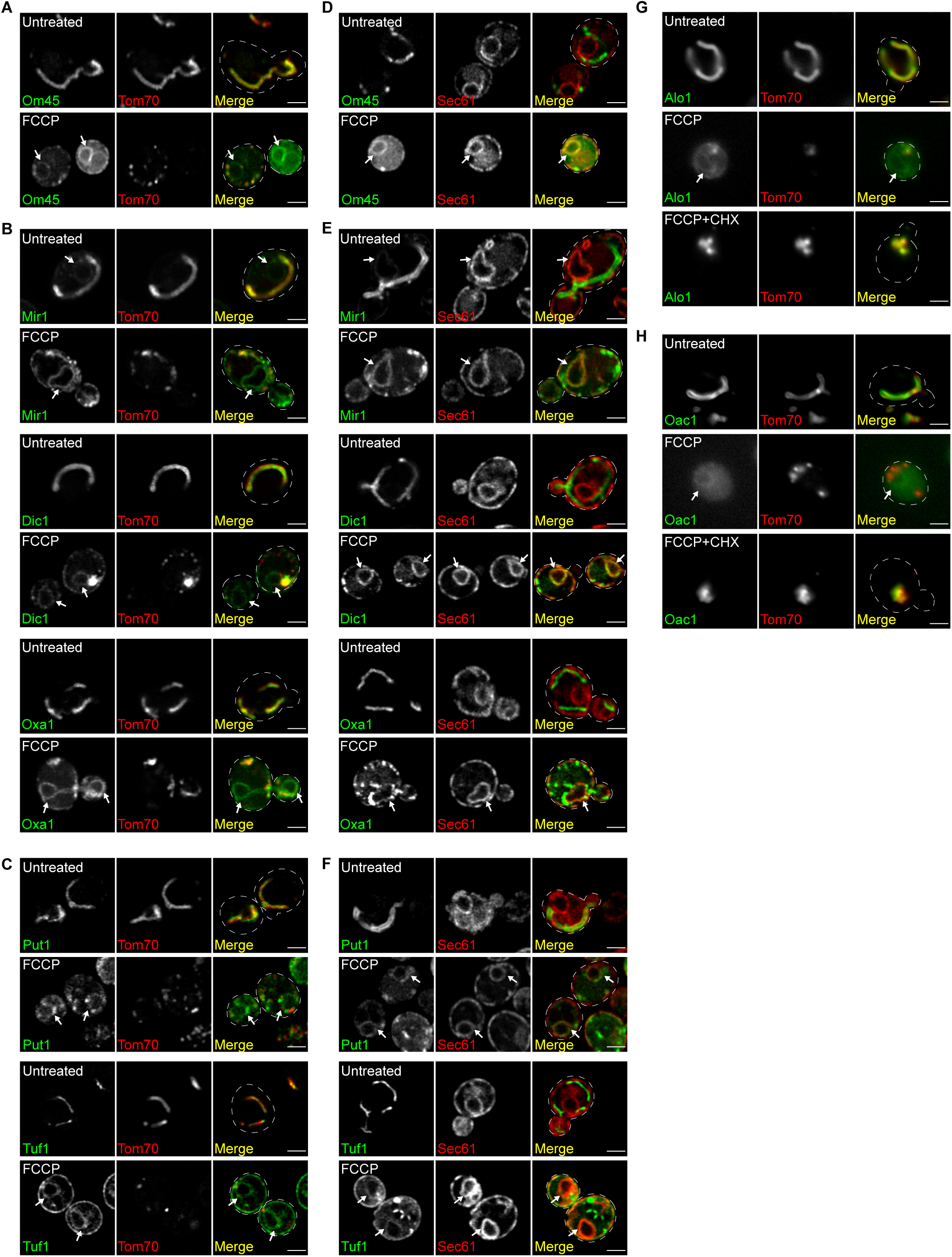
**Related to Fig. 1** (A-C) Super-resolution images of yeast expressing indicated OMM proteins (A), IMM proteins (B) or mitochondrial proteins with unknown sub-organelle localization (C) tagged with GFP and Tom70-mCherry -/+ FCCP. Sub-organelle localizations of mitochondrial proteins were obtained from SGD. (D-F) Super-resolution images of yeast expressing indicated OMM proteins (D), IMM proteins (E) or mitochondrial proteins with unknown sub-organelle localization (F) tagged with GFP and Sec61-mCherry -/+ FCCP. (G and H) Widefield images of yeast expressing Alo1-GFP (G) or Oac1-GFP (H) and Tom70-mCherry -/+ FCCP -/+ CHX (cycloheximide). White arrows mark perinuclear ER. Images show single focal plane. Scale bar = 2 μm.

**Figure S2.**
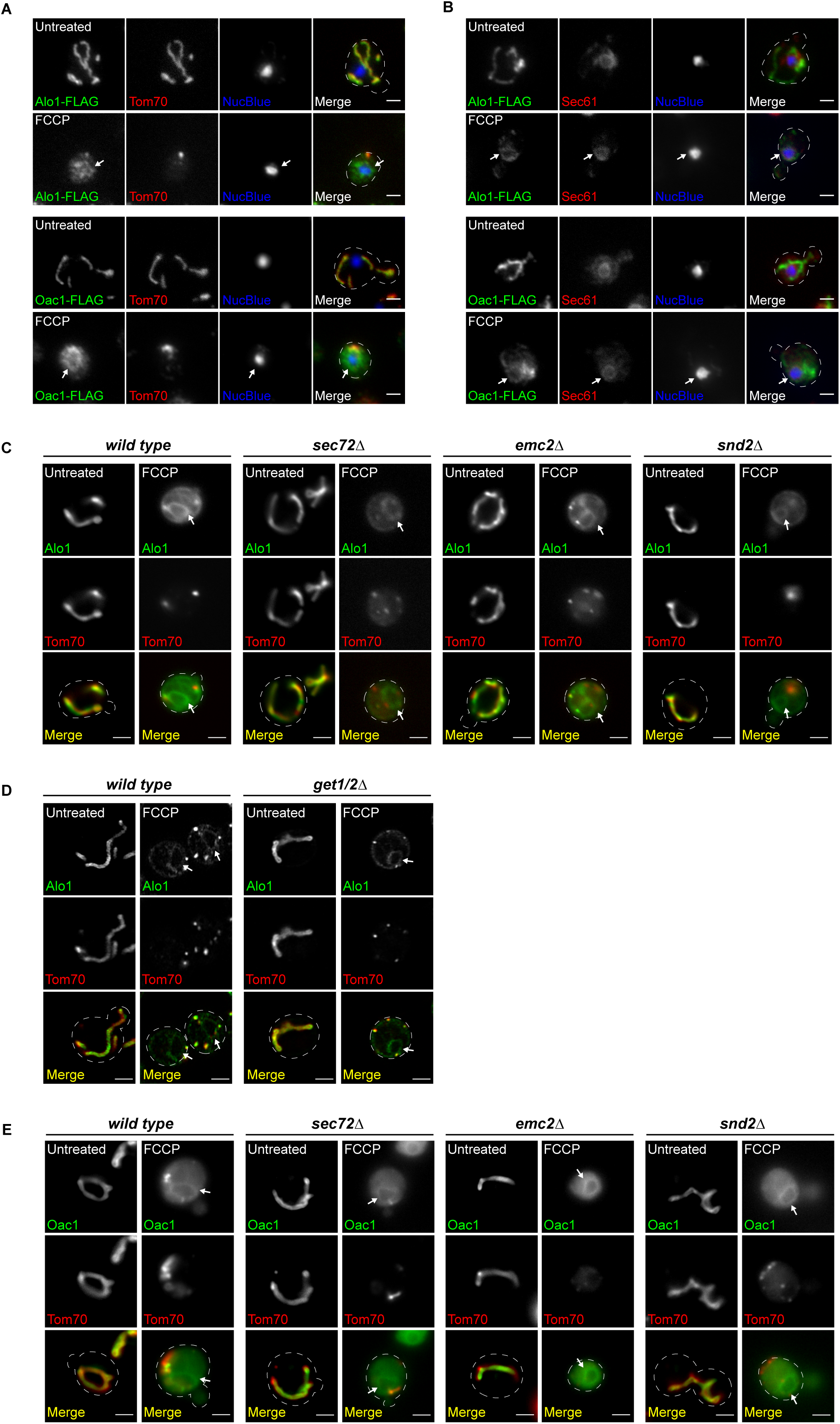
**Related to Fig. 1 and Fig. 2** (A and B) Widefield images of indirect immunofluorescence staining against the FLAG epitope in yeast expressing Alo1- or Oac1-FLAG and Tom70-mCherry (A) or Sec61-mCherry (B) -/+ FCCP. Nucleus stained with NucBlue. (C) Widefield images of wild-type or the indicated mutant yeast expressing Alo1-GFP and Tom70-mCherry -/+ FCCP. (D) Super-resolution images of wild-type or *get1/2*Δ mutant yeast expressing Alo1-GFP and Tom70-mCherry -/+ FCCP. (E) Widefield images of wild-type or the indicated mutant yeast expressing Oac1-GFP and Tom70-mCherry -/+ FCCP. White arrow marks perinuclear ER. Images show single focal plane. Scale bar = 2 μm.

**Figure S3.**
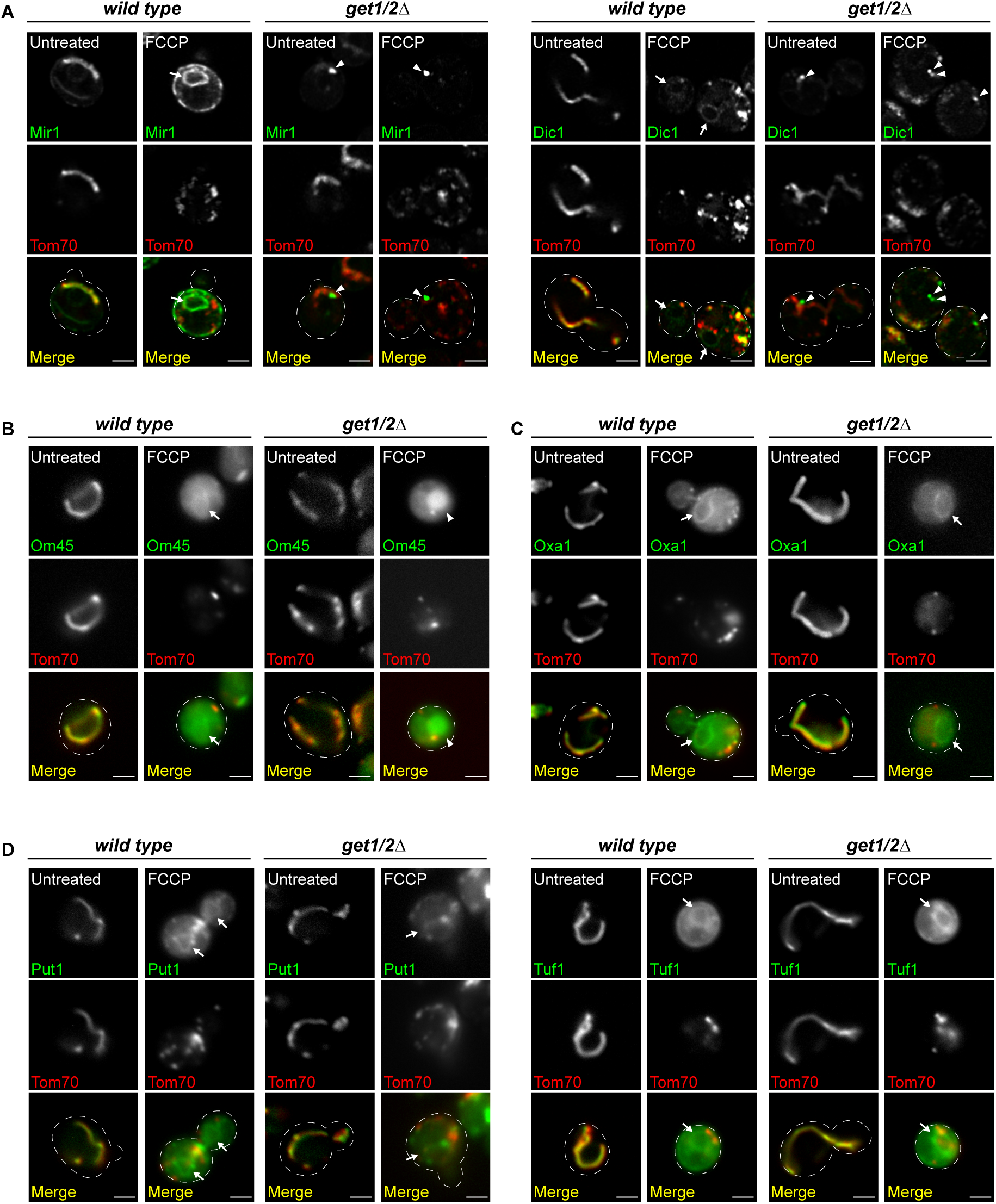
**Related to Fig. 2** (A) Super-resolution images of wild-type or *get1/2*Δ mutant yeast expressing IMM carrier proteins Mir1- or Dic1-GFP and Tom70-mCherry -/+ FCCP. White arrowhead marks protein foci containing GFP-tagged mitochondrial proteins. (B) Widefield images of wild-type or *get1/2*Δ yeast expressing OMM protein Om45-GFP and Tom70-mCherry -/+ FCCP. White arrow marks perinuclear ER. White arrowhead marks vacuole. (C and D) Widefield images of wild-type or *get1/2*Δ mutant yeast expressing IMM protein Oxa1-GFP (C) or mitochondrial matrix protein Put1- or Tuf1-GFP (D) and Tom70-mCherry -/+ FCCP. White arrow marks perinuclear ER. Images show single focal plane. Scale bar = 2 μm.

**Figure S4.**
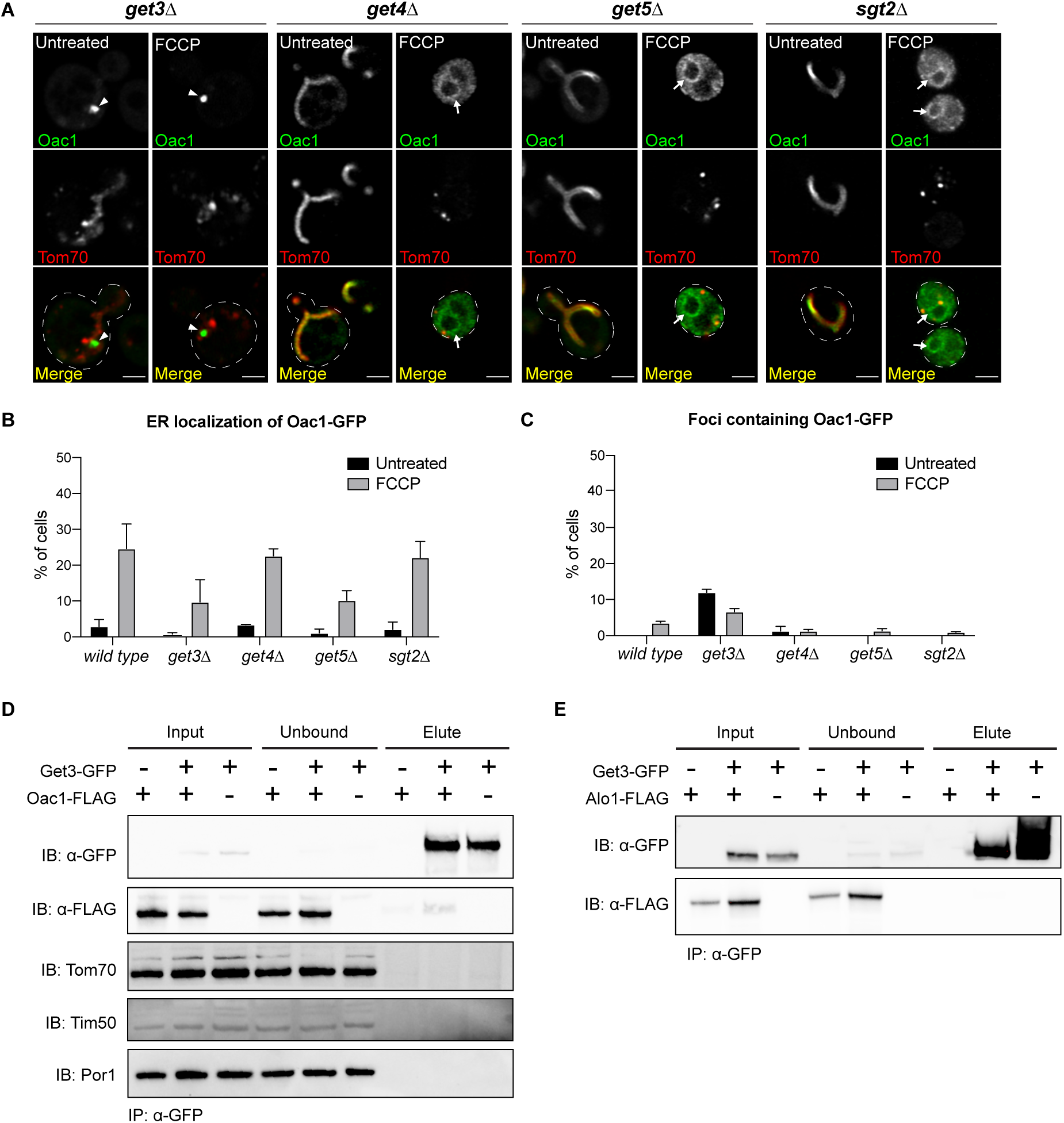
**Related to Fig. 2** (A) Super-resolution images of the indicated mutant yeast expressing Oac1-GFP and Tom70-mCherry -/+ FCCP. White arrows mark perinuclear ER. White arrowhead marks protein foci containing Oac1-GFP. Images show single focal plane. Scale bar = 2 μm. (B and C) Quantification of (A) showing the percentage of cells with Oac1-GFP localized to the ER (B) or cytosolic foci (C) in wild-type or the indicated mutant cells. N > 100 cells per replicate, error bars = SEM of three replicates. (D) Western blot probing for GFP, FLAG-tag, Tom70, Tim50 and Por1 in input, unbound or elution products of immunoprecipitated Get3-GFP in the indicated yeast strains. (E) Western blot probing for GFP and FLAG-tag in input, unbound or elution products of immunoprecipitated Get3-GFP in the indicated yeast.

**Table S1.**
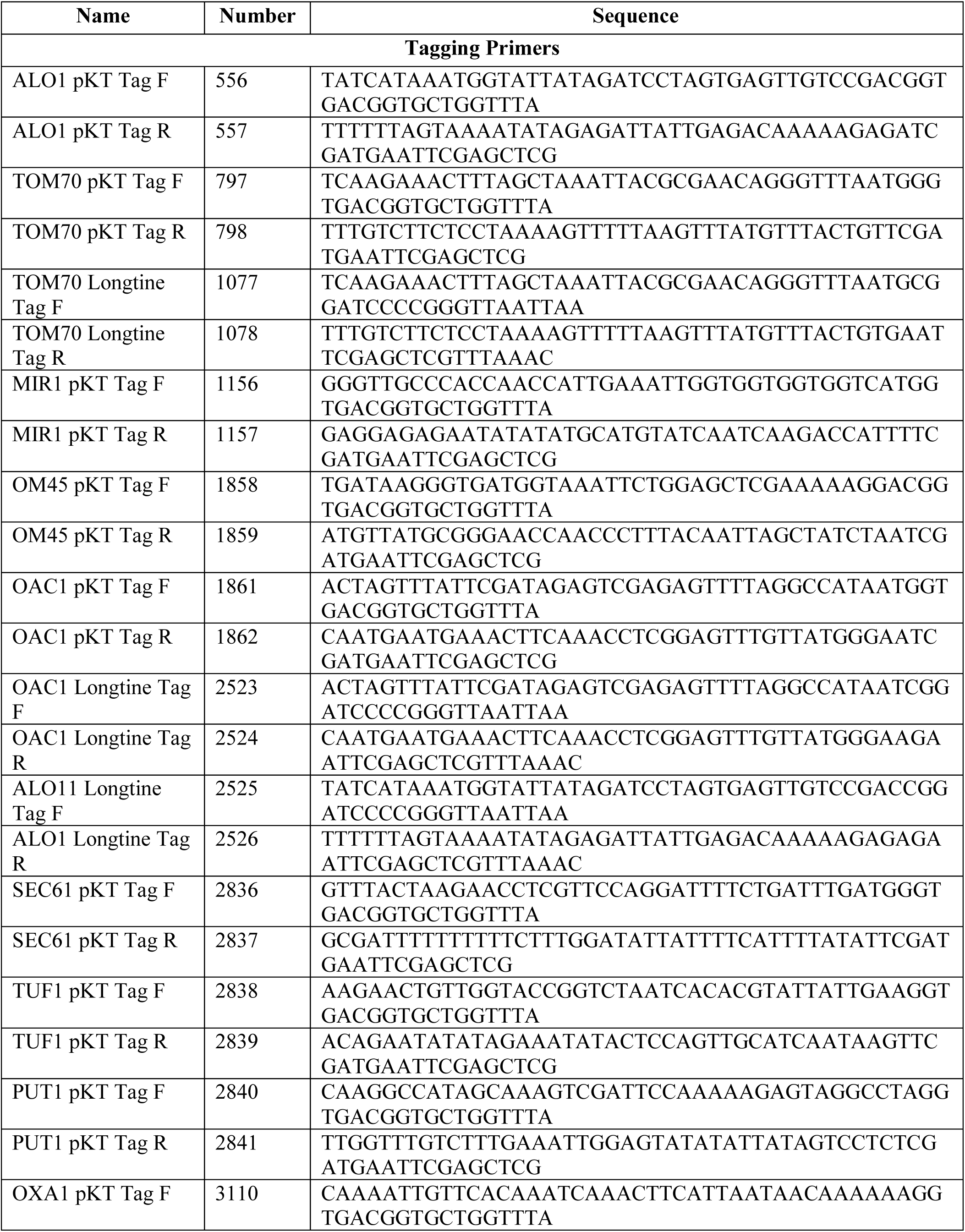
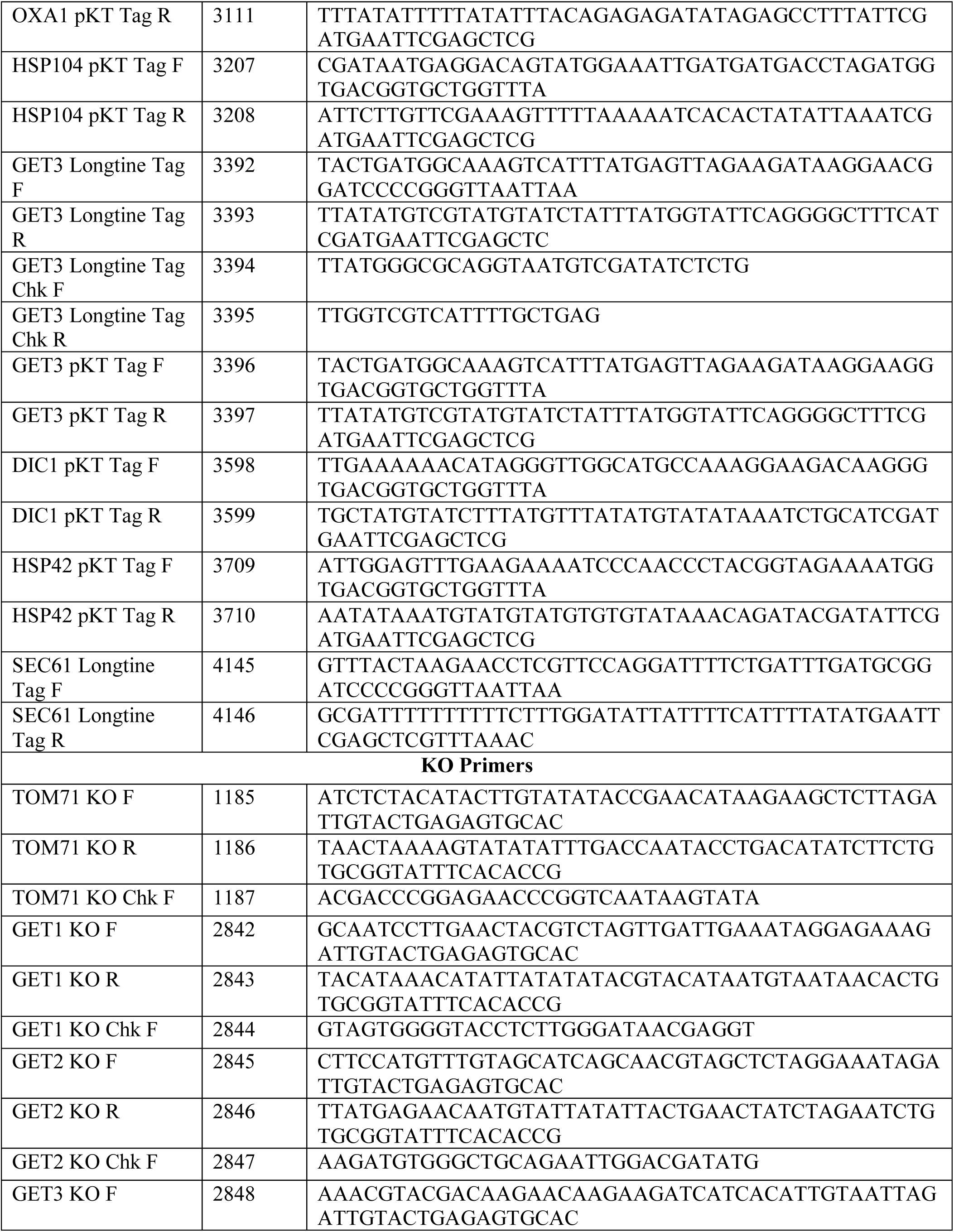
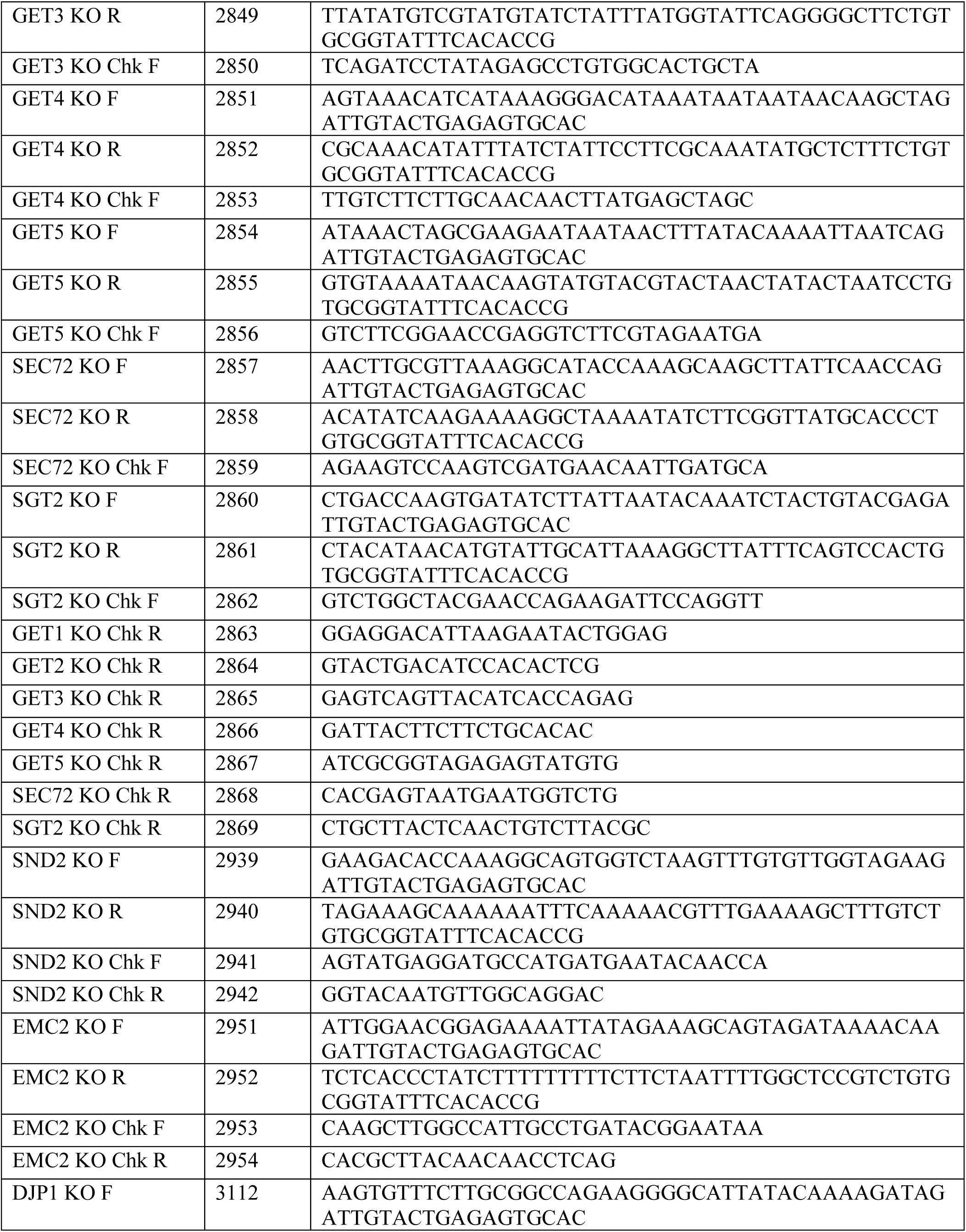
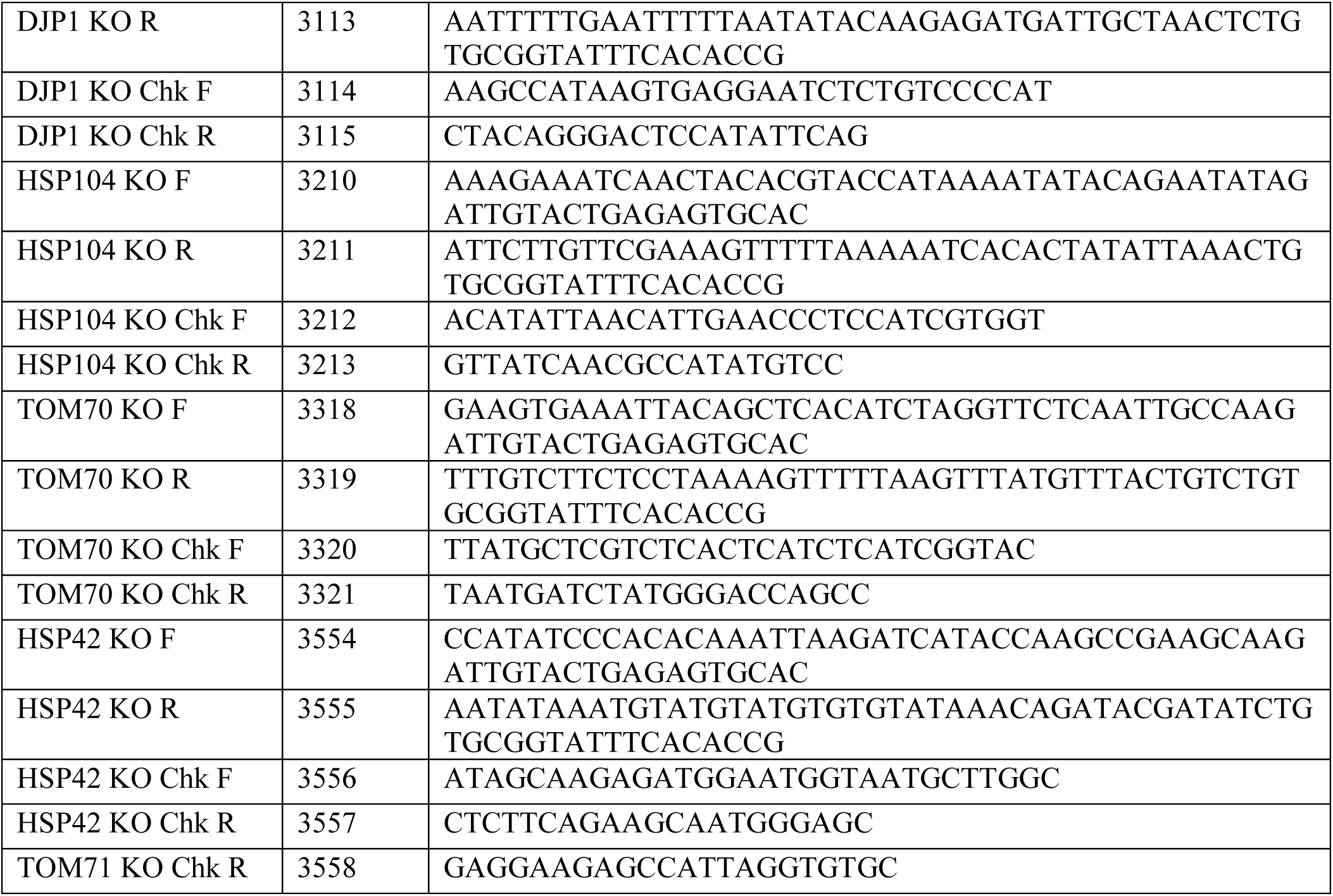
Oligos used in this study

## STAR METHODS

### LEAD CONTACT AND MATERIALS AVAILABILITY

Further information and requests for resources and reagents should be directed to and will be fulfilled by the Lead Contact, Adam L. Hughes. All unique/stable reagents generated in this study are available from the Lead Contact without restriction.

### EXPERIMENTAL MODEL AND SUBJECT DETAILS

#### Yeast Strains

All yeast strains are derivatives of *Saccharomyces cerevisiae* S288c (BY)(Brachmann et al., 1998) and are listed in Supplementary Table 1. Strains expressing tagged proteins from their native loci were created by one step PCR-mediated C-terminal endogenous epitope tagging using standard techniques and the oligo pairs listed in Supplementary Table 2(Brachmann et al., 1998; Sheff and Thorn, 2004). Plasmid templates for GFP tagging were from the pKT series of vectors(Sheff and Thorn, 2004). Plasmid templates for mCherry tagging were from the pKT series of vectors(Sheff and Thorn, 2004) or pFA6a-mCherry-HphMX (Addgene 39295)(Wang et al., 2014b). Integrations were confirmed by correct localized expression of the fluorophore by microscopy. Plasmid template for FLAG tagging was the pFA6a-5FLAG-KanMX6 (Addgene 15983)(Noguchi et al., 2008). Integrations were confirmed by a combination of colony PCR across the chromosomal insertion site and correct band size by western blot. Deletion strains were created by one step PCR-mediated gene replacement using the oligos pairs listed in Supplementary table 2 and plasmid templates from the pRS series vectors(Brachmann et al., 1998). Correct integrations were confirmed with colony PCR across the chromosomal insertion site.

#### Yeast Cell Culture and Media

Yeast cells were grown exponentially for 15-16 hours at 30°C to a final density of 2-7×10^6^ cells/mL prior to starting any treatments. Cells were cultured in YPAD medium (1% yeast extract, 2% peptone, 0.005% adenine, 2% glucose). For FCCP treatment, overnight log-phase cell cultures were grown in the presence of FCCP (final concentration of 10 μM) or cycloheximide (100 μg/mL) for 4-5 hours.

### METHOD DETAILS

#### Microscopy

Optical z-sections of live yeast cells were acquired with a ZEISS Axio Imager M2 equipped with a ZEISS Axiocam 506 monochromatic camera, 100x oil-immersion objective (plan apochromat, NA 1.4), a AxioObserver 7 (Carl Zeiss) equipped with a PCO Edge 4.2LT Monochrome, Air Cooled, USB 3 CCD camera with a Solid-State Colibri 7 LED illuminator and 100X oil-immersion objective (Carl Zeiss, Plan Apochromat, NA 1.4), a ZEISS LSM800 equipped with an Airyscan detector, 63x oil-immersion objective (plan apochromat, NA 1.4) or a ZEISS LSM880 equipped with an Airyscan detector, 63x oil-immersion objective (plan apochromat, NA 1.4). Widefield images were acquired with ZEN (Carl Zeiss) and processed with Fiji(Schindelin et al., 2012). Super-resolution images were acquired with ZEN (Carl Zeiss) and processed using the automated Airyscan processing algorithm in ZEN (Carl Zeiss) and Fiji. Individual channels of all images were minimally adjusted in Fiji to match the fluorescence intensities between channels for better visualization. Line scan analysis was performed on non-adjusted, single z-sections in Fiji. All images shown in Figures represent a single optical section.

#### Serial-Dilution Growth Assays

Five-fold serial dilutions of exponentially growing yeast cells were diluted in ddH_2_O and 3 μL of each dilution was spotted onto YPD (1% yeast extract, 2% peptone, 2% glucose). Final concentration of FCCP is 7 μM. Total cells plated in each dilution spot were 5000, 1000, 20, 40, and 8. Plates were cultured at 30°C for 36 hours before obtaining images.

#### Immunoprecipitation and Western Blotting

Immunoprecipitation and western blot were carried out as described previously(Hughes et al., 2016). Cells were grown as described above. 1.2 x 10^8^ total cells were harvested, resuspended in 500 μL of Lysis Buffer (50 mM Tris pH7.5, 150 mM NaCl, 1 mM EDTA, 10% Glycerol, 1% IGEPAL (NP-40 substitute), 100 μM PMSF) and lysed with glass beads using an Omni Bead Ruptor 12 Homogenizer (8 cycles of 20 seconds each). Cells lysates were cleared by centrifugation at 2,000xg for 3 minutes to remove cell debris, followed by centrifugation at 11,000xg for 5 minutes. Supernatant was collected to a new tube. Pellets were resuspended in 50 μL of SUME buffer (1% SDS, 8 M Urea, 10 mM MOPS, pH 6.8, 10 mM EDTA and 10 mM NEM) and heated at 42 °C for 5 minutes. After centrifugation at 11,000xg for 5 minutes, supernatant of cell pellet resuspension was combined with supernatant from lysate clearance centrifugation, and total volume was adjusted to 1 mL by adding lysis buffer. Lysates were incubated with 25 μL of pre-balanced anti-GFP bead slurry (GTMA, GFP-Trap®_MA, chromotek) at 4 °C overnight and then washed 4 times for 10 minutes in lysis buffer.

Immunoprecipitated proteins were eluted by incubating beads in 2x Laemmli Buffer (63 mM Tris pH 6.8, 2% (w/v) SDS, 10% (v/v) glycerol, 1 mg/mL bromophenol blue, 1% (v/v) β-mercaptoethanol) at 90 °C for 10 minutes. Cells extracts and elution products were resolved on Bolt 4-12% Bis-Tris Plus Gels (NW04125BOX, Thermo Fisher) with NuPAGE MES SDS Running Buffer (NP0002-02, Thermo Fisher) and transferred to nitrocellulose membranes. Membranes were blocked and probed in blocking buffer (1x PBS, 0.05% Tween 20, 5% non-fat dry milk) using the primary antibodies for FLAG (ThermoFisher) and GFP (Sigma Millipore) and HRP conjugated secondary antibodies (715-035-150, Jackson Immunoresearch). Blots were developed with SuperSignal West Pico Chemiluminescent substrate (34580, Thermo Fisher) and exposed with a Bio-Rad Chemidoc MP system.

#### Yeast Indirect Immunofluorescence (IIF) Staining

For IIF staining, overnight log-phase cell cultures were grown with or without FCCP for 3.5 hours in YPAD to OD=0.4. Cells were harvested by centrifugation and fixed in 10 mL fixation medium (4% Polyformaldehyde in YPAD) for one hour. Fixed yeast cells were washed with Wash Buffer (0.1 M Tris, pH=8, 1.2 M Sorbitol) twice and incubated in 2 mL DTT Buffer (10 mM DTT in 0.1 M Tris, pH=9.4) at room temperature for 10 minutes.

Spheroplasts were generated by incubating cells in 2 mL Zymolyase Buffer (0.1 M KPi, pH=6.5, 1.2 M Sorbitol, 0.25 mg/mL Zymolyase) at 30°C for 30 minutes. Spheroplasts were gently diluted in 1:40 using Wash Buffer and attached to glass slides pre-coated with 0.1% poly-L-Lysine (2 mg/mL). Samples were permeabilized in cold 0.1% Triton-X100 in PBS for 10 minutes at 4 °C, briefly dried and blocked in Wash Buffer containing 1% BSA at room temperature for 30 minutes. After blocking, samples were incubated with primary antibody (Monoclonal ANTI-FLAG® M2 antibody produced in mouse, 1:200 diluted in Wash Buffer containing 1% BSA) for 1.5 hours at room temperature and secondary antibody (Goat anti-Mouse IgG (H+L) Cross-Adsorbed Secondary Antibody, Alexa Fluor 488, 1:300 diluted in Wash Buffer containing 1% BSA) for 45 minutes at room temperature. Samples were washed 10 times after each incubation with Wash Buffer containing 1% BSA and 0.1% Tween-20. Slides were washed twice with Wash Buffer before sealing, and mounted with hardset medium (ProLong™ Glass Antifade Mountant with NucBlue™ Stain (P36981), Invitrogen) overnight. Widefield images were acquired as described above.

### QUANTIFICATION AND STATISTICAL ANALYSIS

The number of replicates, what *n* represents, and dispersion and precision measures are indicated in the figure legends. In general, quantifications show the mean ± standard error from three biological replicates with *n* = 100 cells per experiment. In experiments with data depicted from a single biological replicate, the experiment was repeated with the same results.

### DATA AND CODE AVAILABILITY

This study did not generate datasets or code.

## KEY RESOURCES TABLE

| REAGENT or RESOURCE | SOURCE | IDENTIFIER |
| --- | --- | --- |
| **Antibodies** |  |  |
| DYKDDDDK Tag Polyclonal Antibody | Invitrogen | Cat # PA1-984B<br>RRID: AB_347227 |
| Anti-GFP from mouse IgG1κ | Sigma-Aldrich | Cat # 11814460001<br>RRID: AB_390913 |
| Monoclonal ANTI-FLAG® M2 antibody produced in mouse | Sigma-Aldrich | Cat # F1804<br>RRID: AB_262044 |
| Rabbit polyclonal anti-Tom70 | Dr. Nikolaus Pfanner | N/A |
| Rabbit polyclonal anti-Tim50 | Dr. Nikolaus Pfanner | N/A |
| Porin Monoclonal Antibody (16G9E6BC4) | Invitrogen | Cat # 459500<br>RRID: AB_2532239 |
| Peroxidase AffiniPure Donkey Anti-Mouse IgG (H+L) | Jackson Immunoresearch | Cat # 715-035-150<br>RRID: AB_2340770 |
| Goat anti-Mouse IgG (H+L) Cross-Adsorbed Secondary Antibody, Alexa Fluor 488 | Invitrogen | Cat # A-11001<br>RRID: AB_2534069 |
| **Chemicals, Peptides, and Recombinant Proteins** |  |  |
| Carbonyl cyanide 4-(trifluoromethoxy)phenylhydrazone (FCCP) | Sigma-Aldrich | Cat # C2920; CAS # 370-86-5 |
| Cycloheximide | Sigma-Aldrich | Cat # C1988; CAS # 66-81-9 |
| cOmplete Protease Inhibitor Cocktail | Sigma-Aldrich | Cat # 11697498001 |
| Phenylmethylsulfonyl fluoride | Sigma-Aldrich | Cat # P7626; CAS # 329-98-6 |
| IGEPAL® | Sigma-Aldrich | Cat # CA-630; CAS # 9002-93-1 |
| Zymolyase 100T | US Biological Life Sciences | Cat # Z1004; CAS # 37340-57-1 |
| Triton X-100 | Bio-Rad | Cat # 1610407; CAS # 9002-93-1 |
| Formaldehyde 16% in aqueous solution, EM Grade | VWR | Cat # 100503-914; CAS # 50-00-0 |
| DTT (Dithiothreitol) (> 99% pure) Protease free | GOLDBIO | Cat # DTT10; CAS # 27565-41-9 / 3483-12-3 |
| **Experimental Models: Organisms/Strains** |  |  |
| BY4741 MATa his3Δ1 leu2Δ0 ura3Δ0 met15Δ0 | Brachman *et al.*, 1998; ATCC | Cat # 201388 |
| BY4743 MATa/MATα his3Δ1/his3Δ1 leu2Δ0/leu2Δ0 ura3Δ0/ura3Δ0 met15Δ0/+ lys2Δ0/+ | Brachman *et al.*, 1998; ATCC | Cat # 201390 |
| BY4741 TOM70-mCherry:KanMX ALO1-yeGFP:HisMX | This study | AHY3352 |
| BY4741 TOM70-mCherry:KanMX OM45-yeGFP:HisMX | This study | AHY3933 |
| BY4741 TOM70-mCherry:KanMX TUF1-yeGFP:HisMX | This study | AHY6859 |
| BY4741 TOM70-mCherry:KanMX TUF1-yeGFP:HisMX | This study | AHY6860 |
| BY4741 TOM70-mCherry:KanMX PUT1-yeGFP:HisMX | This study | AHY6861 |
| BY4741 SEC61-mCherry:KanMX ALO1-yeGFP:HisMX | This study | AHY6950 |
| BY4741 TOM70-mCherry:KanMX ALO1-yeGFP:HisMX sec72Δ::URA3 | This study | AHY7061 |
| BY4741 TOM70-mCherry:KanMX MIR1-yeGFP:HisMX get1Δ::URA3 get2Δ:HygMX | This study | AHY7110 |
| BY4741 TOM70-mCherry:KanMX ALO1-yeGFP:HisMX get1Δ::URA3 get2Δ::HygMX | This study | AHY7112 |
| BY4741 SEC61-mCherry:KanMX OM45-yeGFP:HisMX | This study | AHY7143 |
| BY4741 SEC61-mCherry:KanMX OAC1-yeGFP:HisMX | This study | AHY7147 |
| BY4741 SEC61-mCherry:KanMX PUT1-yeGFP:HisMX | This study | AHY7149 |
| BY4741 SEC61-mCherry:KanMX TUF1-yeGFP:HisMX | This study | AHY7151 |
| BY4743 TOM70-mCherry:KanMX/+ MIR1-yeGFP:HisMX/+ | This study | AHY7219 |
| BY4741 TOM70-mCherry:KanMX TUF1-yeGFP:HisMX get1Δ::URA3 get2Δ::HygMX | This study | AHY7238 |
| BY4741 TOM70-mCherry:KanMX OM45-yeGFP:HisMX get1Δ::URA3 get2Δ::HygMX | This study | AHY7244 |
| BY4741 TOM70-mCherry:KanMX OAC1-yeGFP:HisMX | This study | AHY7296 |
| BY4741 TOM70-mCherry:KanMX OAC1-yeGFP:HisMX | This study | AHY7297 |
| BY4741 TOM70-mCherry:KanMX PUT1-yeGFP:HisMX get1Δ::URA3 get2Δ::HygMX | This study | AHY7389 |
| BY4741 TOM70-mCherry:KanMX OAC1-yeGFP:HisMX get1Δ::URA3 get2Δ::HygMX | This study | AHY7391 |
| BY4743 SEC61-mCherry:KanMX/+ MIR1-yeGFP:HisMX/+ | This study | AHY7552 |
| BY4741 SEC61-mCherry:KanMX OAC1-yeGFP:HisMX tom70Δ::URA3 tom71Δ::LEU2 | This study | AHY7612 |
| BY4741 SEC61-mCherry:KanMX ALO1-yeGFP:HisMX tom70Δ::URA3 tom71Δ::LEU2 | This study | AHY7616 |
| BY4741 TOM70-mCherry:KanMX ALO1-yeGFP:HisMX snd2Δ::URA3 | This study | AHY7776 |
| BY4741 TOM70-mCherry:KanMX ALO1-yeGFP:HisMX emc2Δ::URA3 | This study | AHY7781 |
| BY4741 TOM70-mCherry:KanMX OAC1-yeGFP:HisMX djp1Δ::URA3 | This study | AHY8186 |
| BY4741 GET3-mCherry:KanMX OAC1-yeGFP:HisMX get1Δ::URA3 get2Δ::HygMX | This study | AHY8389 |
| BY4741 Δ HSP104-mCherry:KanMX OAC1-yeGFP:HisMX get1Δ::URA3 get2Δ::HygMX | This study | AHY8391 |
| BY4741 TOM70-mCherry:KanMX OAC1-yeGFP:HisMX get4Δ::URA3 | This study | AHY8418 |
| BY4741 TOM70-mCherry:KanMX OAC1-yeGFP:HisMX get5Δ::URA3 | This study | AHY8420 |
| BY4741 TOM70-mCherry:KanMX OAC1-yeGFP:HisMX get3Δ::URA3 | This study | AHY8426 |
| BY4741 TOM70-mCherry:KanMX OAC1-yeGFP:HisMX sgt2Δ::URA3 | This study | AHY8428 |
| BY4741 GET3-5FLAG:KanMX | This study | AHY8541 |
| BY4741 TOM70-mCherry:KanMX OAC1-yeGFP:HisMX get1Δ::URA3 get2Δ::HygMX hsp104Δ::NatMX | This study | AHY8629 |
| BY4741 GET3-5FLAG:KanMX OAC1-yeGFP:HisMX | This study | AHY8754 |
| BY4741 OAC1-5FLAG:KanMX | This study | AHY8756 |
| BY4741 OAC1-5FLAG:KanMX GET3-yeGFP:HisMX | This study | AHY8867 |
| BY4741 GET3-yeGFP:HisMX | This study | AHY8941 |
| BY4741 OAC1-yeGFP:HisMX | This study | AHY8943 |
| BY4741 TOM70-mCherry:KanMX OXA1-yeGFP:HisMX get1Δ::URA3 get2Δ::HygMX | This study | AHY9139 |
| BY4741 TOM70-mCherry:KanMX DIC1-yeGFP:HisMX get1Δ::URA3 get2Δ::HygMX | This study | AHY9141 |
| BY4741 TOM70-mCherry:KanMX OAC1-yeGFP:HisMX snd2Δ::URA3 | This study | AHY9149 |
| BY4741 TOM70-mCherry:KanMX OXA1-yeGFP:HisMX | This study | AHY9234 |
| BY4741 TOM70-mCherry:KanMX DIC1-yeGFP:HisMX | This study | AHY9236 |
| BY4741 HSP42-mCherry:KanMX OAC1-yeGFP:HisMX get1Δ::URA3 get2Δ::HygMX | This study | AHY9378 |
| BY4741 TOM70-mCherry:KanMX OAC1-yeGFP:HisMX get1Δ::URA3 get2Δ::HygMX djp1Δ::NatMX | This study | AHY9411 |
| BY4741 get3Δ::HygMX | This study | AHY9417 |
| BY4741 TOM70-mCherry:KanMX OAC1-yeGFP:HisMX get1Δ::URA3 get2Δ::HygMX hsp42Δ::NatMX | This study | AHY9595 |
| MATa his3Δ leu2Δ ura3Δ met15Δ tom70Δ::HygMX | This study | AHY9732 |
| MATα his3Δ leu2Δ ura3Δ lys2Δ tom70Δ::HygMX | This study | AHY9733 |
| MATa his3Δ leu2Δ ura3Δ met15Δ tom71Δ::NatMX | This study | AHY9734 |
| MATα his3Δ leu2Δ ura3Δ lys2Δ tom71Δ::NatMX | This study | AHY9735 |
| MATa his3Δ leu2Δ ura3Δ met15Δ tom70Δ::HygMX tom71Δ::NatMX | This study | AHY9736 |
| MATα his3Δ leu2Δ ura3Δ lys2Δ tom70Δ::HygMX tom71Δ::NatMX | This study | AHY9737 |
| MATα his3Δ leu2Δ ura3Δ lys2Δ get1Δ::KanMX | This study | AHY9754 |
| MATa his3Δ leu2Δ ura3Δ met15Δ get2Δ::KanMX | This study | AHY9755 |
| MATα his3Δ leu2Δ ura3Δ lys2Δ tom70Δ::HygMX get1Δ::KanMX | This study | AHY9759 |
| MATa his3Δ leu2Δ ura3Δ tom70Δ::HygMX get2Δ::KanMX | This study | AHY9760 |
| MATα his3Δ leu2Δ ura3Δ lys2Δ tom71Δ::NatMX get1Δ::KanMX | This study | AHY9765 |
| MATa his3Δ leu2Δ ura3Δ met15Δ tom71Δ::NatMX get2Δ::KanMX | This study | AHY9766 |
| MATα his3Δ leu2Δ ura3Δ tom71Δ::NatMX tom71Δ::NatMX get1Δ::KanMX | This study | AHY9771 |
| MATa his3Δ leu2Δ ura3Δ lys2Δ tom71Δ::NatMX tom71Δ::NatMX get2Δ::KanMX | This study | AHY9772 |
| BY4741 SEC61-mCherry:KanMX OAC1-yeGFP:HisMX get1Δ::URA3 get2Δ::HygMX | This study | AHY10109 |
| BY4741 ALO1-5FLAG:KanMX | This study | AHY10168 |
| BY4741 ALO1-5FLAG:KanMX GET3-yeGFP:HisMX | This study | AHY10170 |
| BY4741 GET3-mCherry:KanMX OAC1-yeGFP:HisMX | This study | AHY10236 |
| BY4741 Δ TOM70-mCherry:KanMX OAC1-yeGFP:HisMX emc2Δ::URA3 | This study | AHY10327 |
| BY4741 Δ TOM70-mCherry:KanMX OAC1-yeGFP:HisMX sec72Δ::URA3 | This study | AHY10329 |
| BY4741 SEC61-mCherry:KanMX OXA1-yeGFP:HisMX | This study | AHY10331 |
| BY4741 SEC61-mCherry:KanMX DIC1-yeGFP:HisMX | This study | AHY10333 |
| BY4741 OAC1-5FLAG:KanMX TOM70-mCherry:HygMX | This study | AHY10410 |
| BY4741 OAC1-5FLAG:KanMX SEC61-mCherry:HygMX | This study | AHY10412 |
| BY4741 ALO1-5FLAG:KanMX TOM70-mCherry:HygMX | This study | AHY10414 |
| BY4741 ALO1-5FLAG:KanMX SEC61-mCherry:HygMX | This study | AHY10416 |
| Oligonucleotides |  |  |
| See Table S1 |  |  |
| Recombinant DNA |  |  |
| Plasmid: pRS40Hyg | Daniel Gottschling | N/A |
| Plasmid: pRS40Nat | Daniel Gottschling | N/A |
| Plasmid: pRS400 | Daniel Gottschling | N/A |
| Plasmid: pRS306 | Sikorski and Hieter, 1989 | N/A |
| Plasmid: pKT128 | Sheff and Thorn, 2004; Addgene | Plasmid # 8729 |
| Plasmid: pKT127-mCherry | Daniel Gottschling | N/A |
| Plasmid: pFA6a-mCherry-HphMX | Wang, 2014; Addgene | Plasmid # 39295 |
| Plasmid: pFA6a-5FLAG-KanMX6 | Noguchi and Garabedian, 2008; Addgene | Plasmid # 15983 |
| Software and Algorithms |  |  |
| FIJI | Schindelin et al., 2012 | Version 1 |
| Prism | GraphPad Software, Inc. | Version 8 |
| SnapGene | GSL Biotech | Version 4.2 |
| ZEN Black Edition | Carl Zeiss Microscopy | Version 2.3 |
| ZEN Blue Edition | Carl Zeiss Microscopy | Version 2.6 |
| Photoshop CC | Adobe | Version 19 |
| Image Lab | Bio-Rad | Version 6 |

